# Cadherin Fat2 directs cellular mechanics to promote epithelial rotation

**DOI:** 10.1101/117515

**Authors:** Ivana Viktorinová, Ian Henry, Pavel Tomancak

**Affiliations:** Max-Planck Institute of Molecular Cell Biology and Genetics, Pfotenhauerstr. 108, 01307 Dresden, Germany; Scionics Computer Innovation GmbH, Löscherstr. 16, 01309 Dresden, Germany

**Keywords:** Fat2, planar cell polarity, collective cell migration, epithelial rotation, left and right symmetry breaking, non-muscle Myosin II, actin, cell mechanics, force generation, myosin pulses, cell shape, cell membrane deformation, tissue morphogenesis

## Abstract

Left and right symmetry breaking is involved in many developmental processes that are important to form bodies and organs. One of them is the epithelial rotation of developing organs. However, how epithelial cells move, how they break symmetry to define common direction of their collective movement and what function rotational epithelial motions have in morphogenesis remain elusive. Here, we identified a dynamic actomyosin network with preferred retrograde contractility at the basal side of the rotating follicle epithelium in *Drosophila* oogenesis. We provide evidence that unidirectional epithelial rotation is a result of actomyosin asymmetry cue transmission onto a tissue plane synchronized by the atypical cadherin Fat2, a key planar cell polarity regulator in *Drosophila* oogenesis. We found that Fat2 directs actomyosin contractility to move the epithelial tissue in order to provide directed elongation of follicle cells. In contrast, loss of Fat2 results in anisotropic non-muscle Myosin II pulses that are disorganized in plane and deform cell shape, tissue and *Drosophila* eggs. Our data indicate that directed elongation of follicle cells is critical for proper *Drosophila* egg morphogenesis. Together, we demonstrate the importance of atypical cadherins in the control of cell mechanics, left/right symmetry breaking and its propagation onto the tissue scale to facilitate proper organ morphogenesis. This process may be evolutionarily conserved in rotating animal organs.

---

Functional organ morphogenesis^1-3^ has been linked to left and right (LR) turns and rotations of epithelial sheets^4 5 6 7 8 9 10^ relative to the organ or body anterior-posterior (AP) axis. A primary determinant of this LR chirality has been associated with the cytoskeleton in different species^11 12 13 14 15^. In rotating *Drosophila* organs such as hindgut^10^ and male genitalia^6^, the consistent handedness of epithelial rotation depends on *myosinID* (*myoID*) and utilizes asymmetric cellular intercalations. Completely different rotational movement was recently identified in the *Drosophila* ovary^8^, where organ-like structures called egg chambers display rotation of an edgeless, monolayered follicle epithelium together with underlying germline cells (called nurse cells and the oocyte), which rotate through the surrounding rigid extracellular matrix (ECM)^8^ (Figure 1a). In contrast to the *Drosophila* hindgut and male genitalia where cell membranes adopt a specific form of asymmetry called planar cell chirality (PCC), the follicle epithelium displays no membrane PCC (Extended Data Fig. 1a, Online Methods) and different egg chamber units in one animal can rotate clockwise or anti-clockwise performing more than three full rotations around their AP axis during early and mid oogenesis^8,16^. This suggests that an alternative, possibly *myoID*-independent, mechanism drives this collective cell behaviour. Interestingly, the basal side of each follicle cell displays clear local chirality of actin-rich protrusions and chiral localization of several planar cell polarity (PCP) molecules genetically implicated in egg chamber rotation^17-20 9 21 22 23^. Epithelial rotation is initially slow during early oogenesis (stages 1-5: average speed ~ 0.2 μm/min)^19^ accelerates in mid oogenesis (stages 6-8: average speed ~ 0.5-0.6 μm/min)^8,9,19,24^ and stops at stage9, ^8^. It has been shown that microtubules (MTs) predict the direction of epithelial rotation in early and mid-oogenesis and their global alignment is regulated by the atypical cadherin Fat2^25 24^. Fat2 is a key PCP regulator of the actin cytoskeleton^25^, basement membrane components^26 20^ and its function is required for epithelial rotation and elongation of *Drosophila* egg chambers^25 9^. The Fat2 asymmetric planar polarized pattern on the basal lagging-membrane side of a follicle cell depends on MTs during fast epithelial rotation^9^. There is no evidence that MTs represent the active force-generating mechanism that drives epithelial rotation, which has been recently shown to involve the actin-rich protrusions^19 23^. However, non-muscle myosin II (Myo-II) that generally provides contractility and force generation to actin cytoskeleton is missing on actin-rich protrusions^19^. Therefore, motivated by the observation that pharmacological depletion of non-muscle myosin II (Myo-II) leads to no epithelial rotation^9^, we hypothesized that the basal actin filaments that contain Myo-II are better candidates to fulfill the force generating function. To test this hypothesis, we investigated the function of Myo-II, its connection to the PCP pathway in *Drosophila* epithelial rotation and the role of their interplay in egg chamber morphogenesis.

**FIGURE 1.**
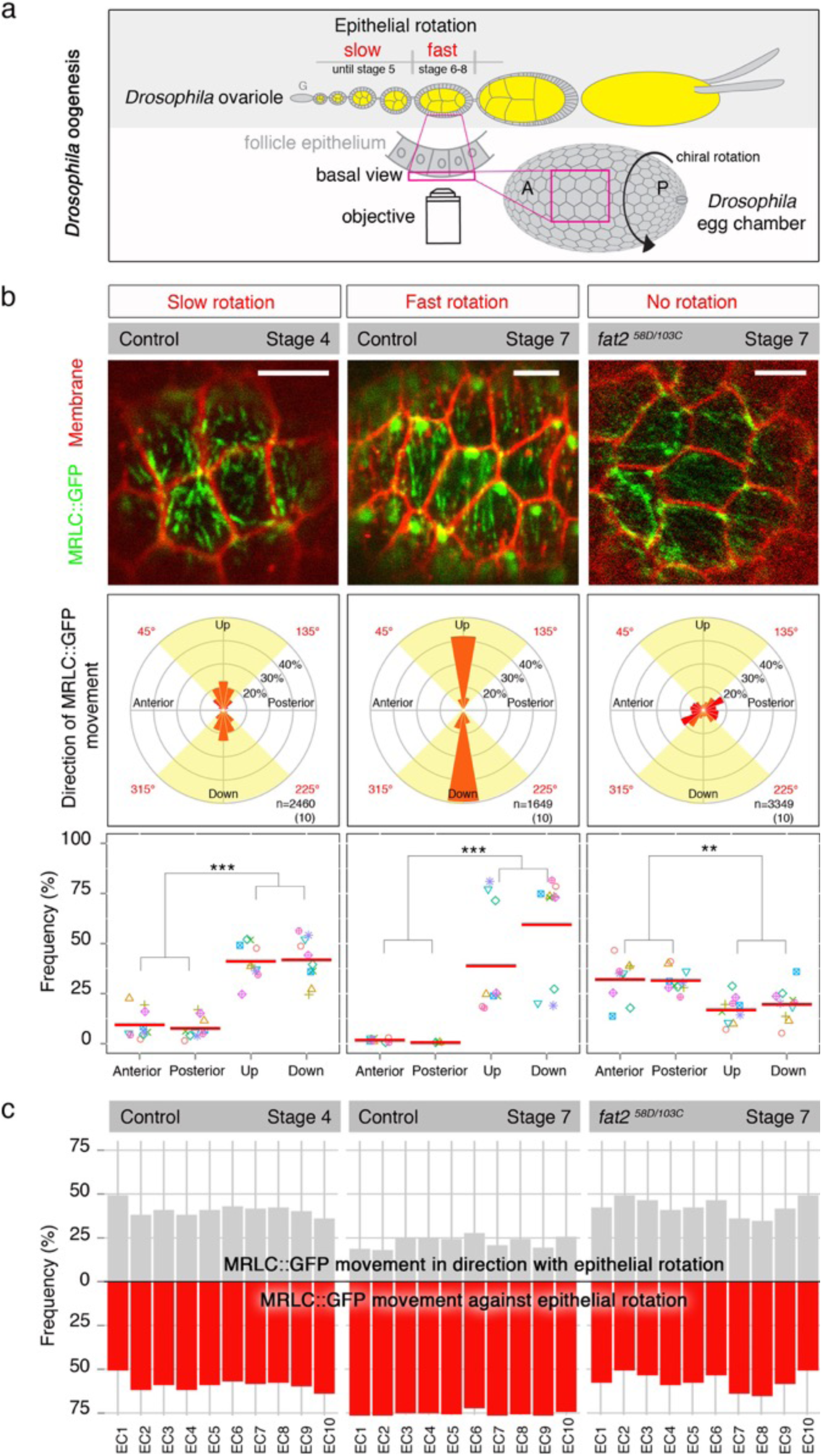
Global Myo-II retrograde movement defines direction of epithelial rotation in the Fat2-dependent manner. **(a) *Drosophila*** ovaries consist of ovarioles, which contain egg chambers of different stages. They bud from the germarium (G) and undergo initially slow (clockwise or anti-clockwise, i. e. chiral) epithelial rotation (until stage 5), which accelerates during stages 6-8 and ceases at stage 9. The confocal view of analyzed basal side of the follicle epithelium, covering nurse cells and the oocyte (yellow), is indicated. A = anterior, P = posterior. **(b)** First row: Basal MRLC::GFP localization (green) at the basal side of the follicle epithelium during slow (stage 4), fast (stage 7) and no epithelial rotation (***fat2*** mutant stage 7) of ***Drosophila*** egg chambers. Scale bars = 5μm. Anterior is on the left. Second row: Angular distribution of MRLC::GFP movement is shown in 20 degree-bin rose diagrams. The numbers of MRLC::GFP individual signals and egg chambers (in brackets) analyzed is at the lower right. 0° represents MRLC::GFP movements towards anterior. The number on the circle represents the fraction of MRLC::GFP signal movements in a bin as a percentage of all MRLC::GFP movements analyzed. Rose diagram indicates four 45-degree quadrants: Anterior (315<45°), Posterior (135<225°), Up (45<135°, yellow) and Down (225<315°, yellow). Third row: Fractions of MRLC::GFP movement within an image frame over time (Online Methods) are shown for indicated quadrants and individual egg chambers (displayed by different color and shape). Note that MRLC::GFP moves preferentially in Up and Down quadrants during slow and fast rotation and this is lost in ***fat2*** mutant egg chambers. Mean (red bar) and *P*-values < 0.001 (^***^) and <0.01 (^**^) are shown. **(c)** Frequencies of MRLC::GFP movement of Up (grey) and Down (red) quadrants for 10 individual egg chambers (EC) when the direction of the epithelial rotation is unified towards up. Note that the preferred direction of MRLC::GFP movement is towards down (against epithelial rotation), i.e. retrograde. MRLC::GFP movements in ***fat2*** mutant egg chambers were artificially unified (Online Methods).

## RESULTS

### Highly dynamic Myo-II behaviour at the basal side of the follicle epithelium

In order to understand Myo-II function in epithelial rotation, we first investigated the behaviour of the Myo-II regulatory light chain (MRLC, called Spaghetti Squash, *sqh* in *Drosophila*) fused to GFP (MRLC::GFP) in a null *sqh^AX3^* mutant^27^ at the basal cortex of the follicle epithelium using *ex vivo* live imaging. We analyzed Myo-II behaviour in three different situations: slow (stage 4), fast (stage 7) and no epithelial rotation (stage 7 of *fat2^58D/103C^* mutants in a null *sqh^AX3^* background since *fat2* mutant egg chambers display no epithelial rotation^9,19^) (Extended Data Fig. 1b). High-speed confocal live imaging of MRLC::GFP uncovered a very dynamic pattern of Myo-II in a thin layer (≤1000 nm) at the basal cortex of the follicle epithelium during slow, fast and no rotation (Figure 1b and Movies 1, 2, 3). We distinguished individual MRLC::GFP dot-like signals with an average size of 363 nm ± 0.05 nm (n = 136) and an average speed of 2.12 μm/min ± 0.8 (n = 101) for slow, 2.44 μm/min ± 0.96 (n = 105) for fast and 1.99 μm/min ± 0.62 (n = 100) for no epithelial rotation. This speed was consistent with the speed of anterograde flow of actomyosin during zebrafish gastrulation^28^. We also observed large intense MLRC::GFP dots (1.01um ± 0.14 um, n= 50, Figure 1b and Movie 2) close to the lagging end of migrating follicle cells, which were lost in the *fat2* mutant follicle epithelium (Figure 1b and Movie 3), suggesting an unknown function in epithelial rotation. Taken together, we discovered highly dynamic behaviour of Myo-II at the basal cortex of the *Drosophila* follicle epithelium.

### Global actomyosin retrograde movement is regulated by atypical cadherin Fat2 in the follicle epithelium

To find out whether the small (~360nm) MRLC::GFP dots moved in a specific direction with respect to the egg chamber axis, we quantified directions of MRLC::GFP movement expressed as angles ranging from 0° to 360° where 0° represented the anterior and 180° the posterior of egg chambers (Extended Data Fig. 1c and Online Methods). In contrast to the situation with no epithelial rotation, which showed no clear preference in direction of MRLC::GFP movement (Figure 1b), we observed that during slow rotation, MRLC::GFP showed weak preferred movement perpendicularly to the AP axis of egg chambers (Figure 1b) that was strongly reinforced during fast rotation (Figure 1b and Extended Data Fig. 1d). Similarly, labeling actin filaments with a LifeAct^29^ molecule fused to GFP (LifeAct::GFP) showed strong preference in LifeAct::GFP movement perpendicularly to the AP axis of egg chamber during fast rotation (Extended Data Fig. 1d). This preference of small MRLC::GFP dots to move perpendicularly to the AP axis of egg chambers as well as the loss of their preferred direction in *fat2* mutant egg chambers observed in live imaging was corroborated by the analysis of MRLC::GFP signal in fixed wild type and *fat2* mutant egg chambers during early and mid-oogenesis (Extended Data Fig. 2).

Having established the global trend of Myo-II movement, we next asked whether individual MRLC::GFP dots moved randomly along their preferred direction (perpendicular to the AP axis of egg chambers) during slow and fast epithelial rotation. To this end, we calculated how frequently MRLC::GFP dots moved within defined angle range of four (90 degrees) quadrants: Anterior (315°≤45°), Up (45°≤135°), Posterior (135°≤225°) and Down (225°≤315°). The data revealed preferred movement within Up and Down quadrants and an asymmetry in movement of MRLC::GFP dots within these quadrants in individual egg chambers (Figure 1b). This asymmetry was initially small during slow rotation and became prominent during fast rotation. In contrast, rather weak to no asymmetry has been detected in *fat2* mutant egg chambers (Figure 1b).

Next we asked in what way this MRLC::GFP movement asymmetry relates to the direction of epithelial rotation. In order to define the average percentage of MRLC::GFP dots moving with or against epithelial rotation, we unified the direction of epithelial rotations in the direction Up for all analyzed egg chambers and detected that on average 59% and 77% MRLC::GFP dots moved against epithelial rotation during slow and fast rotation, respectively (Figure 1c and Extended Data Fig. 3a). This was not true for the *fat2* mutant egg chambers (Figure 1c and Extended Data Fig. 3a), where no preferred global direction was identified in individual egg chambers. We also observed that actin molecules preferably moved (78%) against fast epithelial rotation, based on LifeAct-GFP, and this movement was comparable to the MRLC::GFP movement during fast rotation (Extended Data Fig. 3b and Figure 1c). Thus, we discovered that MRLC::GFP dots preferred to move against epithelial rotation during slow and fast rotation.

Contractile actomyosin rings, Myo-II pulses and Myo-II asymmetric localization on the cell membrane represent features that have been previously linked to tissue movements during epithelial morphogenesis^28 30 31 6^. However, we observed neither of these, indicating that known mechanisms of actomyosin mediated collective epithelial movement do not play a role in this system.

Taken together, our results revealed novel temporally regulated and spatially coordinated Fat2-depenent actomyosin movement at the basal side of the follicle epithelium in the direction opposite to epithelial rotation (retrograde movement).

### Fat2 regulates actomyosin retrograde movement in individual follicle cells

Next, we sought to uncover how Fat2 coordinates retrograde Myo-II movement to break the Myo-II symmetry in the follicle epithelium in order to define the direction of epithelial rotation around the AP axis of egg chambers and how it increases Myo-II retrograde movement during the onset of fast epithelial rotation (Figure 1c). We have shown here that Fat2 regulates global alignment of Myo-II perpendicularly to the AP axis of egg chambers based on live imaging and fixed tissue (Figure 1 and Extended Data Fig. 2). We additionally observed that Fat2 is not required for local (intracellular) alignment of MRLC::GFP in individual follicle cells of fixed egg chambers during slow and fast epithelial rotation (Extended Data Fig. 2). Based on this, we hypothesized that artificial global alignment of individual *fat2* mutant follicle cells perpendicularly to the AP axis of egg chambers should be in principle sufficient to break the symmetry in the follicle epithelium. To simulate such situation, we developed a computational angular correction approach (Figure 2a and Online Methods), in which we assumed that such epithelial remodeling (either by dissolving/re-establishment of adherens junctions or cell intrinsic reorientation of actomyosin cytoskeleton) happens in the wild-type situation with minimal movement of the respective components (i.e. cells or cytoskeleton would rotate 60 rather than 120 degrees to align). To mimic this, individual *fat2* mutant follicle cells were angularly corrected for their main MRLC::GFP direction within the smallest possible angle (i.e. ≤ 90°) to reach perpendicular alignment to the AP axis of the individual egg chambers (Figure 2a and Online Methods).

**FIGURE 2.**
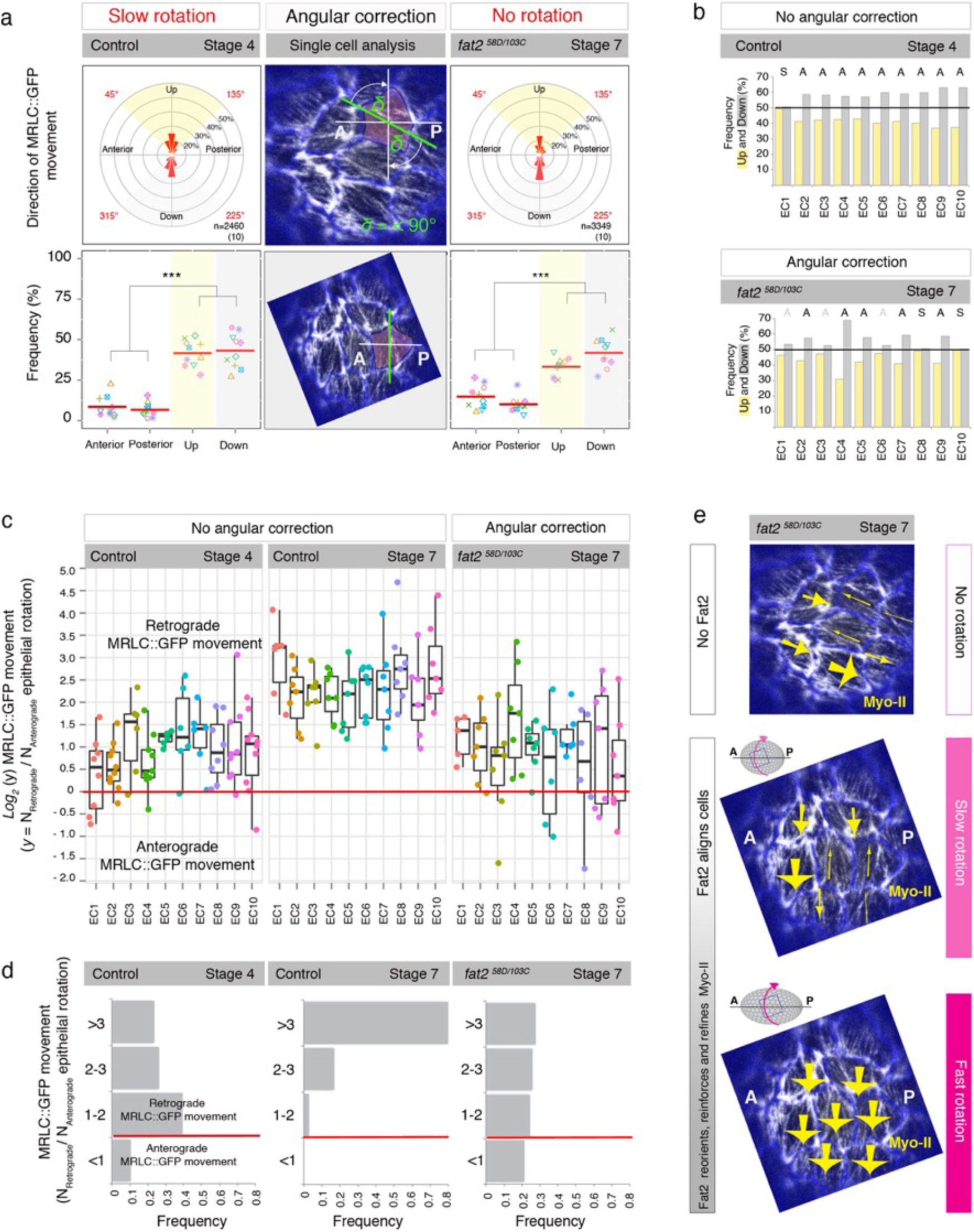
Fat2 globally aligns Myo-II asymmetry to break its global symmetry and locally reorients, refines and reinforces Myo-II in individual follicle cells to promote epithelial rotation. **(a)** First row: Angular distribution and fractions of MRLC::GFP movements as described (Figure 1) for the angularly corrected (Online Methods) stage 4 (slow epithelial rotation) and ***fat2*** mutant stage 7 (no epithelial rotation). Second row: Frequencies of MRLC::GFP movements in four quadrants (see description in Figure 1) and *P*-values < 0.001 (^***^) are shown. Middle images (time projected MRLC::GFP) show example of the angularly corrected image of ***fat2*** mutant follicle cells. A = anterior, P = posterior. **(b)** Frequencies of MRLC::GFP movement in Up and Down quadrants in control stage 4 and the angularly corrected ***fat2*** mutant stage 7 for 10 individual egg chambers. A = asymmetry (black = strong, grey = weak), S = symmetry. **(c)** Weighted ratios of MRLC::GFP moving against (retrograde) or with (anterograde) the direction of epithelial rotation plotted on ***log2*** scale. Dots represent individual follicle cells and color individual egg chambers (EC). Box plots with median for ***log2*** ratio values are shown. Symmetry border is at 0. Note that MRLC::GFP prefers to move mainly in retrograde direction in individual follicle cells but a few of them also showed anterograde direction during weak and no epithelial rotation **(d)** in contrast to significantly unified MRLC::GFP retrograde movement during fast epithelial rotation. **(e)** Model of Fat2 function to globally align and locally reinforce, refine and reorient Myo-II. Note that Myo-II asymmetry is not visible in images and can be revealed only upon Myo-II quantification over time. Yellow arrow indicates direction and the magnitude of the Myo-II asymmetry detected in a follicle cell. As often opposing Myo-II asymmetries were identified in the ***fat2*** mutant follicle cells, our data provide a novel view that neighboring follicle cells cannot sense each other's cytoskeletal direction in ***fat2*** mutant egg chambers as previously thought based on fixed tissues and 0°-180° range^40^.

When angular correction was applied to our data obtained from live imaging, MRLC::GFP dots tended to move perpendicularly to the AP axis in the *fat2* mutant follicle epithelia of stage 7 (Figure 2a), however, only to the extent observed in the angularly corrected control stage 4 (slow epithelial rotation) and far less when compared to the original control stage 7 (fast epithelial rotation, no angular correction in Figure 1b). This indicated an additional Fat2-dependent mechanism that is required to increase number of MRLC::GFP dots moving perpendicularly to the AP axis of egg chambers. In addition, as fixed follicle cells at control stage 4 did not display proper perpendicular alignment of the MRLC::GFP pattern to the AP axis of egg chambers (Extended Data Fig. 2), we expected an increase in the movement of MRLC::GFP dots perpendicularly to the AP axis of living egg chambers after the angular correction, however, instead we observed that angular corrected control stage 4 did not change (Figure 2a) compared to the original situation (Figure 1b), indicating that the movement of MRLC::GFP dots needed to be additionally refined with the onset of fast epithelial rotation.

Taken together, this analysis revealed that in addition to the requirement of Fat2 to globally align Myo-II of follicle cells perpendicularly to the AP axis of egg chambers, Fat2 is also required to refine Myo-II alignment and reinforce this directed movement of Myo-II in the follicle epithelium to reach fast epithelial rotation (compare to Figure 1b).

To find out whether the global artificial alignment of *fat2* mutant follicle cells is sufficient to break the Myo-II symmetry in the follicle epithelium, we further calculated frequency of MRLC::GFP dots moving in the preferred directions (quadrants Up and Down) in individual egg chambers (Figure 2b and Online Methods). We found that five out of ten egg chambers would clearly break Myo-II symmetry similarly to the control stage 4, three displayed only slight asymmetry and two remained symmetrical (Figure 2a, b), suggesting that global alignment of Myo-II alone could in theory lead to the Myo-II symmetry breaking in the follicle epithelium, however, to reach the Myo-II asymmetry comparable to the control stage 4, an additional Fat2-dependent mechanism was required in ~50% of the egg chambers.

To further understand why only 50% of egg chamber clearly broke Myo-II symmetry after applying the angular correction (Figure 2b), we analyzed behaviour of MRLC::GFP dots in individual follicle cells of control and *fat2* mutant egg chambers. To this end, we plotted the direction of MRLC::GFP dots as the weighted ratio of those moving against epithelial rotation (retrograde MRLC::GFP dots) versus those moving with it (anterograde MRLC::GFP dots) (Figure 2c). The analysis revealed that individual follicle cells displayed different ratios of retrograde/anterograde MRLC::GFP dots, but on average during slow rotation with the magnitude around 1-2 while during fast rotation with the significantly increased magnitude of 3 or more (Figure 2d and Extended Data Fig. 4a,b). Notably, we identified one or more follicle cells of individual egg chambers with symmetric MRLC::GFP movement or with anterograde direction during slow rotation. This was never the case during fast rotation when all egg chambers displayed only follicle cells with retrograde MRLC::GFP movement (Figure 2c). Moreover, we observed the same, exclusively retrograde movement with LifeAct::GFP in individual follicle cells during fast rotation (Extended Data Fig. 4c,d), indicating that MRLC::GFP movement reliably reflects LifeAct::GFP behaviour. Coming back to our question why only 50% of egg chambers clearly broke the Myo-II symmetry, we found that in the angularly corrected *fat2* mutant follicle cells the magnitude of the retrograde/anterograde ratio resembled those found during slow rotation. Importantly, the *fat2* mutant egg chambers with no clear Myo-II asymmetry (Figure 2b) contained one or more follicle cells with strong preferred anterograde MRLC::GFP movement (Figure 2c) and that occurrence was on average more frequent in *fat2* mutant egg chambers than in the control ones during slow rotation (Figure 2d).

Thus, this data uncovered that next to global Fat2 function to align actomyosin cytoskeleton perpendicularly to the AP axis of egg chambers and to define preferred direction of actomyosin contractility by breaking the Myo-II symmetry, Fat2 functions also within individual follicle cells to reorient the anterograde Myo-II movement and to refine and reinforce retrograde Myo-II in individual follicle cells to promote fast epithelial rotation (Figure 2e). In addition, we point out that the weak Myo-II asymmetries detected in the *fat2* mutant follicle cells (Figure 2c) indicate existence of an alternate, initially Fat2-independent mechanism of the Myo-II symmetry breaking in the follicle cells.

Altogether, our data provide evidence that Fat2 propagates the initial, intracellular actomyosin-based asymmetry onto the tissue level to facilitate epithelial rotation of *Drosophila* egg chambers.

### Fat2-dependent epithelial rotation directs elongation of follicle cells

Because loss of Fat2 leads to static and round egg chambers^9^, we next wished to understand what role Fat2-dependent epithelial rotation plays in egg chamber morphogenesis. Epithelial rotation has been clearly linked to proper PCP of basement membrane and actin filaments, which is required for egg chamber elongation along the AP axis^18,19^. In addition, integrin-based adhesions to ECM can modulate speed of epithelial rotation and impact shape/stretching of follicle cells^18^. Thus, we wondered whether epithelial rotation could define the shape of follicle cells and so measured their roundness (Online Methods). Indeed, we found that follicle cells are on average significantly more elongated during fast rotation than follicle cells in static *fat2* mutant egg chambers and less elongated during slow rotation (Figure 3a). We also pharmacologically perturbed epithelial rotation by using actin-depleting drug (Latrunculin A) and adhesion-depleting drug (CK-666, depleting Arp2/3 complex) that have been shown to stop epithelial rotation^9 19^. Both experiments resulted in static follicle cells and similar round shape of *fat2* mutant follicle cells (Figure 3a and Extended Data Fig. 5a), indicating that epithelial rotation is required for elongation of follicle cells.

**FIGURE 3.**
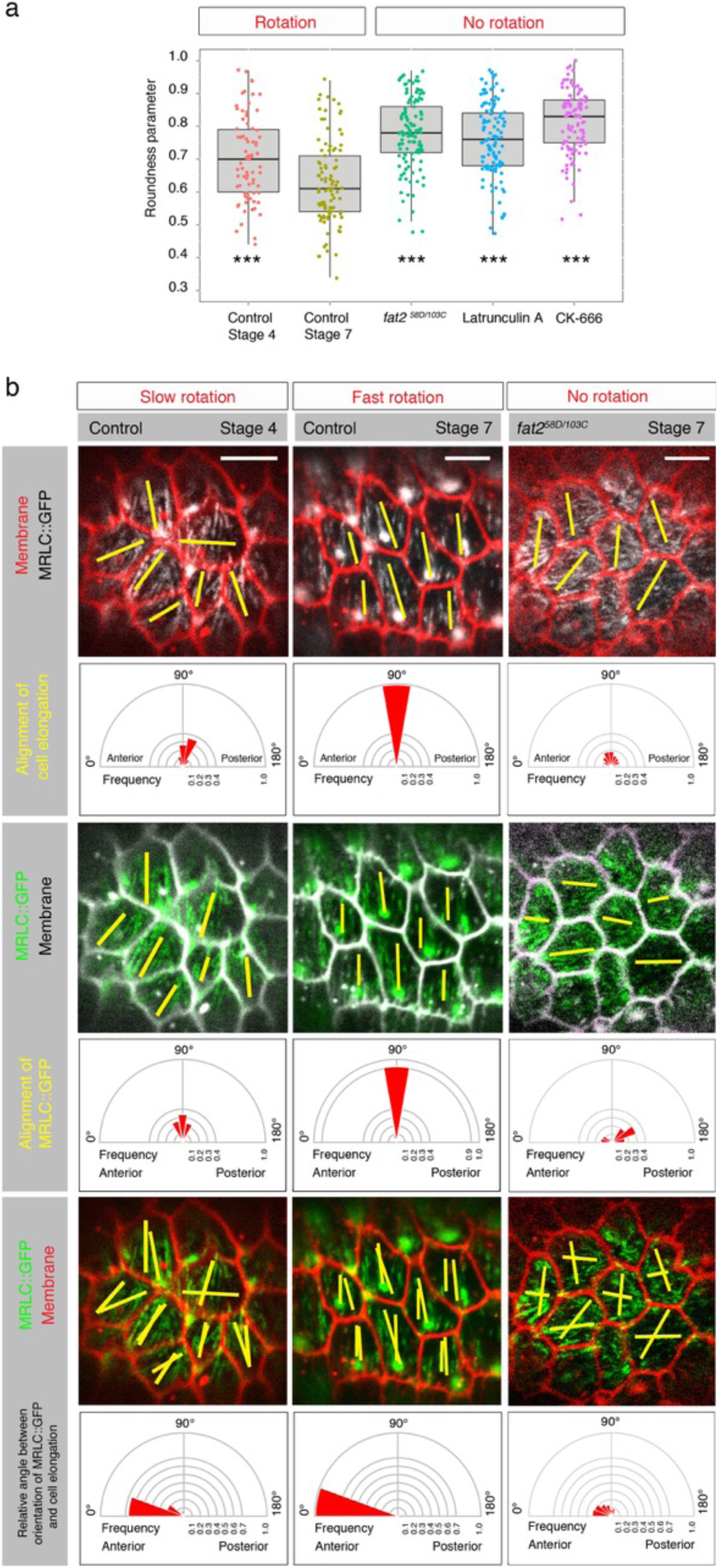
Fat2-dependent epithelial rotation is required for proper elongation of follicle cells. **(a)** Follicle cells were significantly rounder (based on Roundness parameter, Online Methods) in the static follicle epithelium [***fat2*** mutant follicle cells, n = 121 over 10 independent egg chambers; follicle cells depleted for actin, (Latrunculin A), n = 118 over 10 independent egg chambers; follicle cells depleted for cell adhesion (CK-666), n = 107 over 10 individual egg chambers] than in the slow (control stage 4, n = 77 over 9 independent egg chambers) or the fast (control stage 7, n = 97 over 11 individual egg chambers) rotating follicle epithelium. ^***^ = *P*<0.001. **(b)** Angular distribution of alignment of cell elongation and of MRLC::GFP movement expressed as frequencies in 20°-bin rose diagrams and their relative angles are shown for slow (control stage 4), fast (control stage 7) and no (***fat2*** mutant of stage 7) epithelial rotation. Yellow bars show direction on range of 0°-180°. Scale bars = 5μm. Anterior is on the left.

Next, we wished to know whether epithelial rotation could define the direction of elongation of follicle cells. We therefore quantified the alignment of cell elongation expressed as angular direction in 20 degree bins through a range of 0°-180° during slow, fast and no epithelial rotation. We found that follicle cells elongated mainly perpendicularly to the AP axis of egg chambers during fast epithelial rotation, which was weaker during slow rotation (Figure 3b). To understand how the elongation of follicle cells relates to the global alignment of Myo-II (the predicted active force generator together with actin in this tissue) in individual cells, we calculated the relative angle between the global alignment of follicle cell elongation and the MRLC::GFP pattern (Online Methods). We revealed that MRLC::GFP followed follicle cell elongation in 65% during slow epithelial rotation and increased to 100% during fast epithelial rotation (Figure 3b). Interestingly, the global alignment of MRLC::GFP pattern preceded the one of follicle cell elongation during slow epithelial rotation, indicating that epithelial movement prefigures elongation of follicle cells. In contrast, *fat2* mutant follicle cells rather displayed random global alignment of their elongation, the MRLC::GFP pattern and their relative angle.

This data show that Fat2-dependent epithelial rotation is required for proper elongation of follicle cells and the global alignment of their elongation in the direction of epithelial rotation (henceforth called *directed elongation*). This explains why it does not effectively matter whether egg chambers rotate clockwise or anti-clockwise as long as the epithelial rotation is around the AP axis of egg chambers. In addition, supported by the recent findings that shape of follicle cells can determine cellular alignment of MTs in *Drosophila* oogenesis^32^, we further speculate that such directed elongation leads to local and global alignment of MTs at the basal side of follicle cells. Not surprisingly, static egg chambers do not align MTs^9 26^.

### Fat2 suppresses anisotropic and premature Myo-II pulses

We next asked what role directed elongation plays in egg chamber morphogenesis. Interestingly, when we analyzed individual *fat2* mutant follicle cells, besides weak Myo-II asymmetries and rounder cell phenotype, we also observed spatially unequal (anisotropic) MRLC::GFP pulses (Figure 4a, b, c and Extended Data Fig. 5b) with constant remodeling and deformation of cellular membranes, impacting area size (Figure 4d and Extended Data 5b). This pulsatile behaviour was missing in the corresponding control (stage 7) during fast epithelial rotation (Figure 4a). This membrane deformation resulted in significant basal area contractions in *fat2* mutant follicle cells compared to the corresponding control (Figure 4d). However, average area stayed unchanged, indicating that these area contractions were asynchronous among neighboring *fat2* mutant follicle cells (Figure 4d). We observed that the reduction in basal area followed ~6s after the increase of MRLC::GFP (Figure 4e) based on a cross-correlation coefficient ((Figure 4f and Online Methods). Thus, our data provide evidence that Fat2 suppresses anisotropic Myo-II pulses and cellular membrane contractions/relaxations to prevent cell and tissue deformations (Figure 4g). This quantification of live imaging data also extends our previous observation of basally deformed follicle cells in the fixed follicle epithelium of *fat2* mutant egg chambers^25^. Notably, similar physiological Myo-II pulses have been observed during mid-late *Drosophila* oogenesis (starting stage 9 with the peak at stage 10), which have been shown to properly elongate follicle cells and whole egg chambers along their AP axis^33^.

**FIGURE 4.**
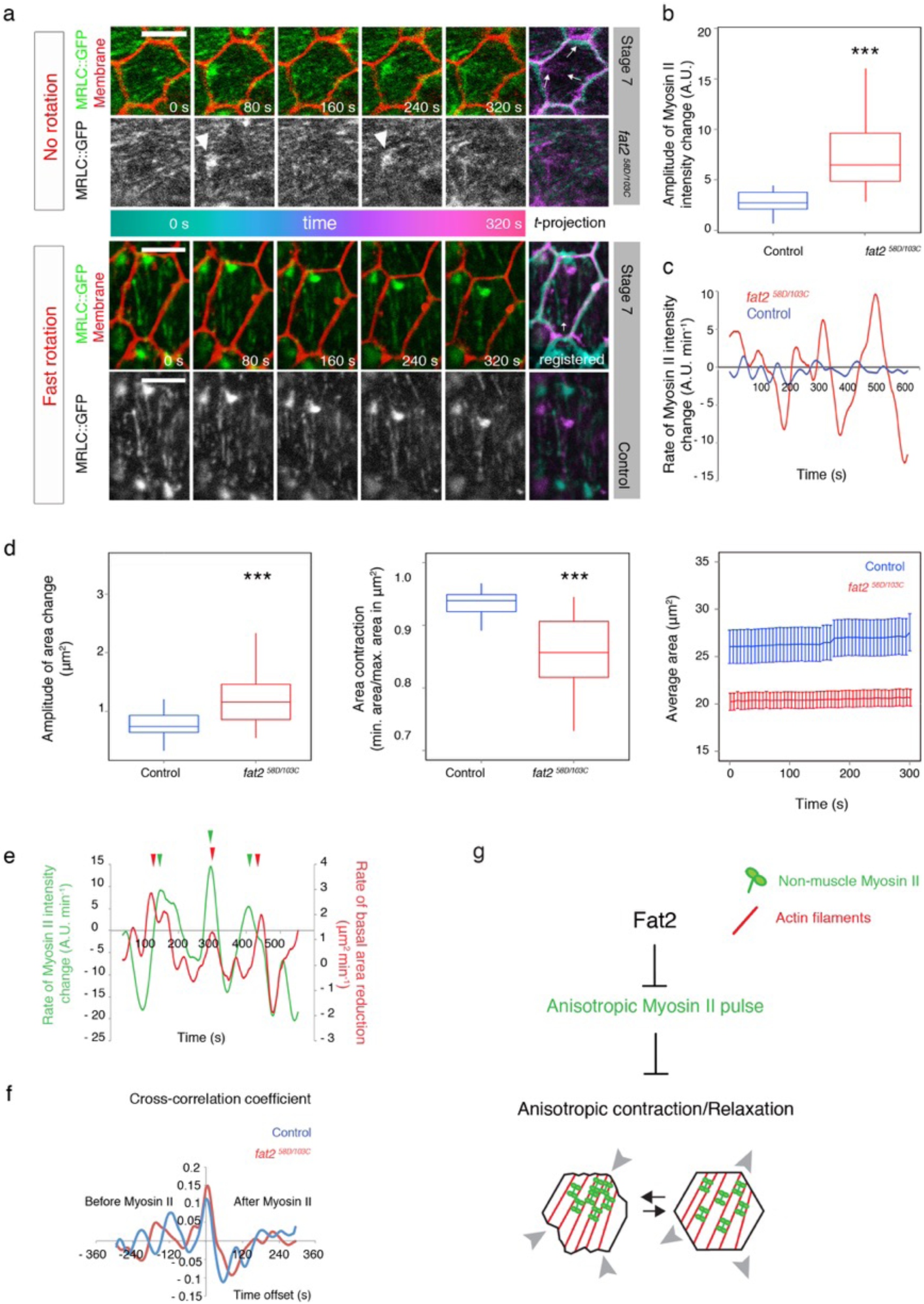
Fat2 guarantees proper follicle cell shape by suppression of premature Myo-II anisotropic pulses. **(a)** Unequal MRLC::GFP intensity increase (Myo-II anisotropic pulse) and membrane deformations are shown in ***fat2*** mutant follicle cells of stage 7. Arrowheads indicate an individual MRLC::GFP intensity increase (a pulse). Note that MRLC::GFP pulses were not observed in control stage 7 (data for one representative cell is shown) **(c)**. Amplitude of Myosin-II intensity change **(b)** and of area change **(d)** were significantly (^***^ = *P*<0.001) higher in ***fat2*** mutant follicle cells of stage 7 than in controls. Correspondingly, area contraction [ratio of the minimal area (contraction) to the maximal area (relaxation)] was significantly weaker in ***fat2*** mutant follicle cells of stage 7. Average area of analyzed follicle cells, which was calculated for ***fat2*** mutant follicle cells of stage 7 (n=56, 7 independent egg chambers) and control stage 7(n=28, 5 independent egg chambers) over time (300s) is shown **(d)**. **(e)** Rate of MRLC::GFP intensity change (green) and rate of the basal area reduction (red) in one representative cell temporarily correlates. Peaks are marked with arrowheads with corresponding color. **(f)** Cross-correlation coefficient (average of all correlation coefficients with different time offsets) between rate of MRLC::GFP intensity and rate of follicle cell area reduction (red line, n=56, 7 independent egg chambers). ***R*** = 0.15 with peak maxima at +6s, indicating that MRLC::GFP precedes the basal area reduction. Blue line shows the average time-dependent correlation of control (wild-type situation of stage 7) between rates of MRLC::GFP intensity change and basal area reduction (n=28, 5 independent egg chambers). ***R*** = 0.113 with a maximum peak at 0, suggesting that MRLC::GFP and basal area reduction are almost simultaneous. **(g)** Schematic model of how Fat2 suppresses unwanted MRLC::GFP anisotropic contractions/relaxations and thereby membrane deformations. Scale bars = 5μm. Anterior is on the left.

In summary, we propose that Fat2 synchronizes the initial default LR Myo-II asymmetry in one direction in the plane (the global Fat2 function), reorients and reinforces actin cytoskeleton within follicle cells (the local Fat2 function), resulting in transmission of actomyosin-based LR asymmetry onto tissue scale. This facilitates epithelial rotation, directed elongation of follicle cells and cell-shape-dependent local and global alignment of MTs. As MTs have been reported to be relevant for turn-over of the Fat2 (zig-zag) asymmetric pattern during fast epithelial rotation^9^, we further hypothesize that Fat2 uses epithelial rotation to align MTs for establishment of its planar polarized asymmetric pattern in a positive-feedback amplification mechanism^9^. This mechanism thus provides a robust Fat2-MTs-dependent platform that guarantees stable global and local alignment of actomyosin and MTs cytoskeleton during fast epithelial rotation (stage 6-8), which upon its establishment and maintenance^9^ likely becomes independent of epithelial rotation as observed in^19^. We envision that this is further critically required for globally aligned, physiological Myo-II pulses at the basal side of follicle cells in order to properly elongate follicle cells and egg chambers along their AP axis in later *Drosophila* oogenesis (Figure 5).

**FIGURE 5.**
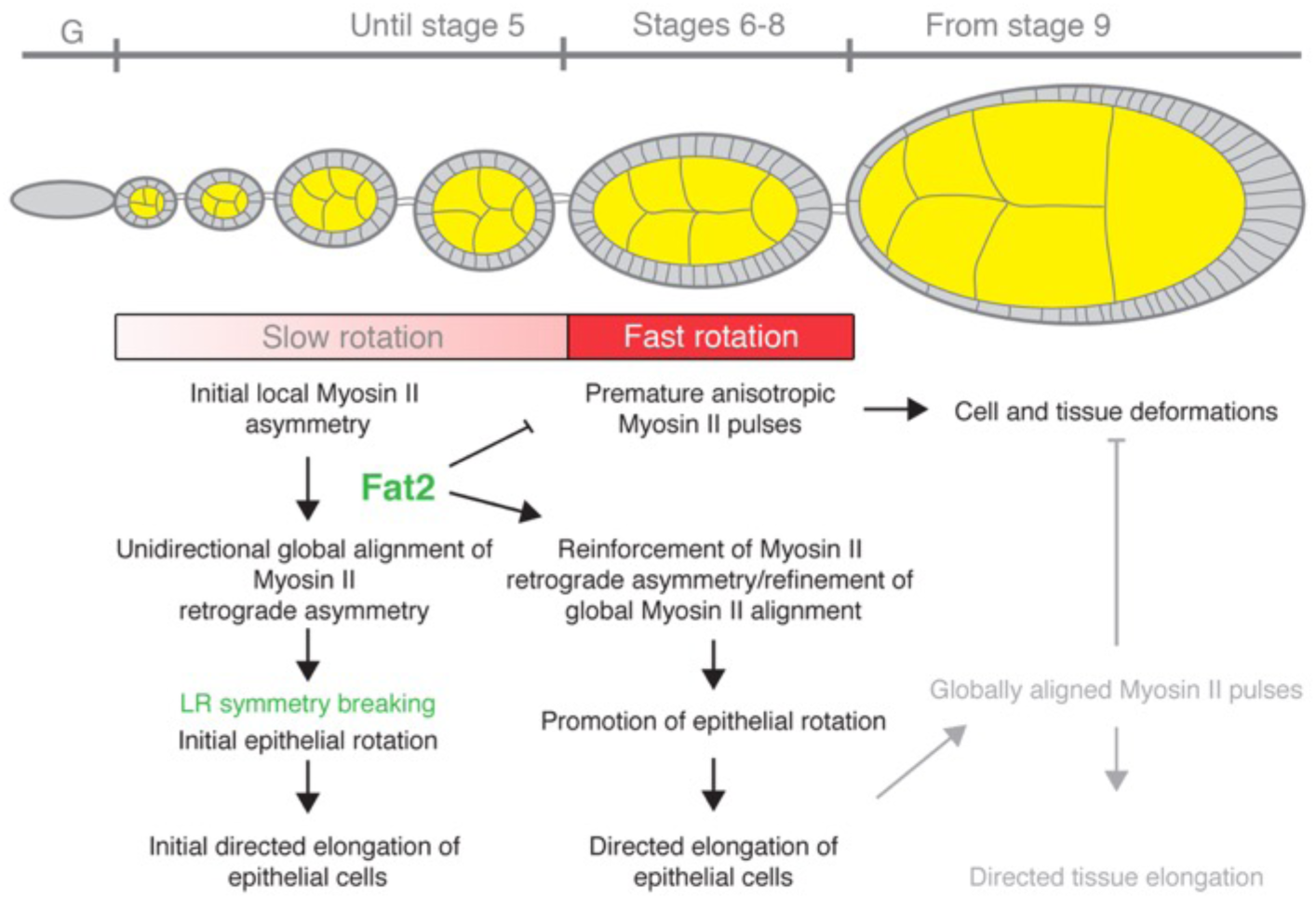
Model how Fat2 breaks left/right symmetry via directed orchestration of actomyosin contractility to promote epithelial rotation and directed elongation of follicle cells. Fat2 globally aligns initial Myo-II asymmetry in individual follicle cells to set up unidirectional global alignment of Myo-II retrograde asymmetry, which breaks left/right symmetry in the early follicle epithelium and initiates epithelial rotation and directed elongation of follicle cells. By stage 6 (start of fast epithelial rotation coinciding with the planar Fat2 asymmetric pattern), Fat2 reinforces Myo-II and refines retrograde asymmetry and reorients anterograde Myo-II in some follicle cells, leading to strong global retrograde Myo-II movement that in turn promotes epithelial rotation and directed elongation of follicle cells. In this respect, Fat2 suppresses anisotropic Myo-II pulses that would otherwise deform follicle cells, the follicle epithelium and egg chambers, resulting in the round phenotype. We propose that the global alignment of cellular cytoskeleton perpendicular to the AP axis of egg chambers is critical for globally aligned Myo-II pulses that oscillates along the dorso-ventral axis of follicle cells later in *Drosophila* oogenesis that elongate follicle cells and egg chambers along their AP axis^33^.

## Discussion

One of the best understood processes that involves actomyosin flow in LR symmetry breaking is single and four cell *C. elegans* embryo^15^, where active torque generates force that leads to LR symmetry breaking and embryo asymmetry. Another example, where actomyosin anterograde and retrograde flow are involved, is the yolk cell during spreading of the enveloping cell layer over it by a process that involves unidirectional movement of strong actomyosin contractile ring in zebrafish embryo^28^. Besides invertebrate and vertebrate embryos^2^, single migrating cells also need to break symmetry to polarize in order to define direction of their motion. For example, actomyosin retrograde flow has been found in non-adhesive cells, allowing pushing of the cell mass forward^34^. In contrast, classical Myosin II-driven actin retrograde flow on the leading edge is used in_single pulling (ECM-adhesive) cells as originally found in neuronal growth cones^35^, however, little is known about dynamics of central stress fibers that retract the majority of the cell mass in direction of cell migration. Here, we identified preferred retrograde Myo-II movement on the basal actin stress-like fibers in ECM-adhesive follicle cells of the rotating *Drosophila* follicle epithelium. We therefore propose directed basal actomyosin contractility a candidate force-generating mechanism that actively moves the cell mass of follicle cells in collective manner and drives epithelial rotation of *Drosophila* egg chambers.

Little is known about the mechanistic control of LR symmetry breaking in tissue morphogenesis. It has been only recently discovered that atypical cadherins can act via force generating molecules such as unconventional myosins Dachs and MyoID^36 37^. Our data show that another atypical cadherin Fat2 can control LR symmetry breaking by regulation of cell mechanics via conventional non-muscle Myo-II. It appears that the atypical cadherin subfamily likely developed a prominent function to shape tissues in two ways dependent on tissue character: i) in migratory tissue via retrograde Myo-II movement and ECM-dependent mechanism (Fat2-Myo-II in *Drosophila* egg chambers in this work) and ii) in moving but non-ECM-migratory tissue via intercalations (Dachsous-MyoID in the *Drosophila* hindgut^37^, Fat-Dachsous-Dachs in the *Drosophila* wing^38^). As Fat2 close homologs, namely Fat1-3, exist in vertebrates^39^, it is likely that similar conserved mechanisms are used to move and sculpt tissues and organs in vertebrates.

## ACKNOWLEDGEMENTS

We thank to Thomas Lecuit for *AX3; Sqh::GFP* fly stock, to Daiki Umetsu for LifeAct::GFP fly stock and to Bloomington Stock Center for the line BL4414. We also thank the Scientific Computing Facility at MPI-CBG (Robert Haase) for developing an in-house Fiji Plugin. We are grateful to Stephan Grill for comments on the manuscript.

## AUTHOR CONTRIBUTIONS

I.V. made the observation of retrograde movement and pulsed contractility, planned the project and performed all experiments. I.V. analyzed all data. I.V. and I.H. analyzed the pulse contractility data together and I. H. helped with the R scripts for data analysis. P.T. generously supported this work. I.V. wrote the manuscript and created figures with help from P.T. and I.H. All authors commented on the manuscript.

## AUTHOR INFORMATION

Reprints and permissions information is available at www.nature.com/reprints. The authors declare no competing financial interests. Correspondence and requests for materials should be addressed to.

## SUPPLEMENTARY INFORMATION

is linked to the online version of the paper at www.nature.com/nature.

---

**Extended Data Figure 1.**
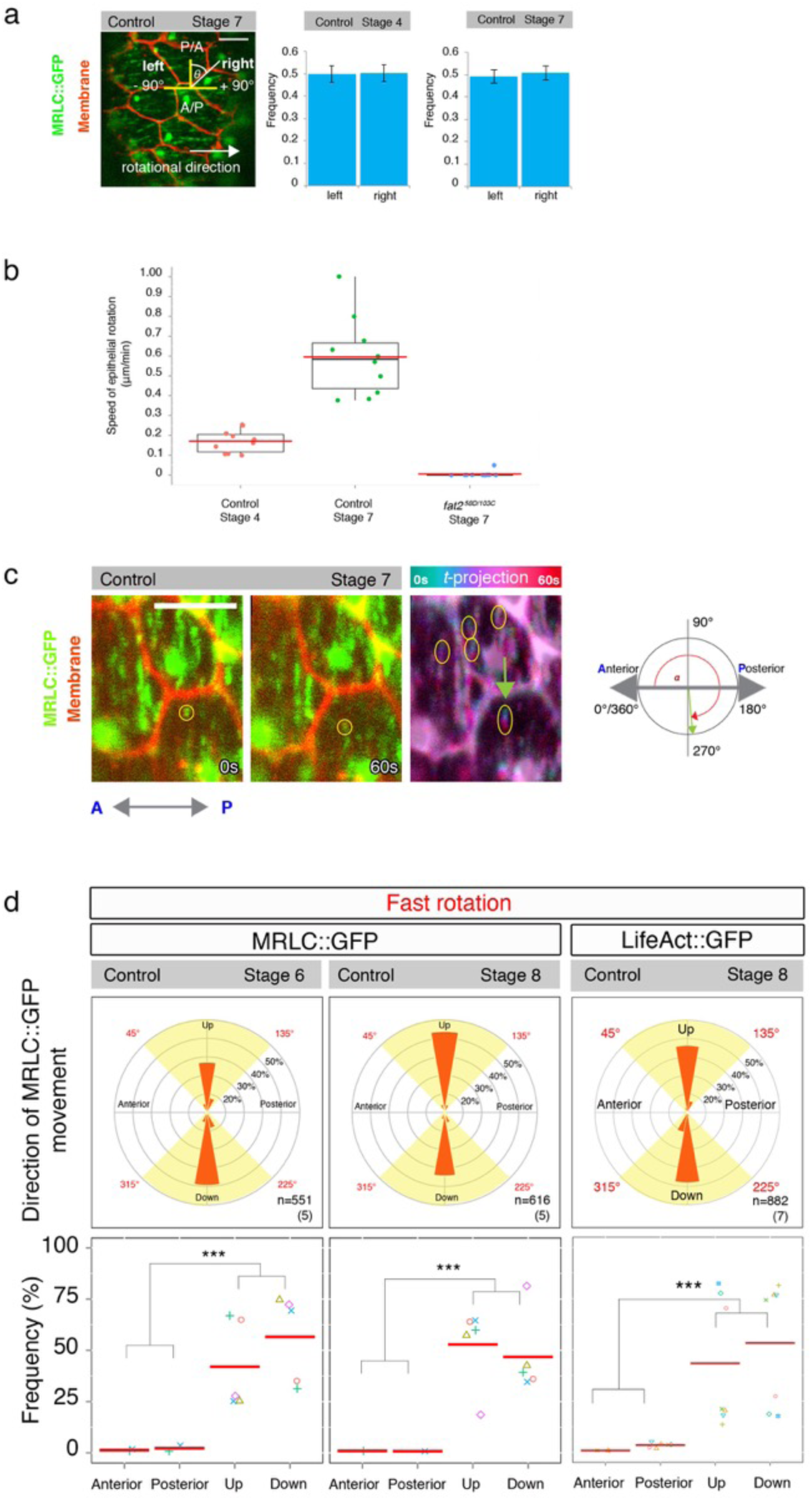
**(a)** Planar cell chirality (PCC) does not contribute to left/right symmetry breaking in the slow (control stage 4) and fast (control stage 7) rotating follicle epithelium. Number of cell membranes n = 261 and n = 319 over 10 independent egg chambers were analyzed for slow (control stage 4) and fast (control stage 7) epithelial rotation, respectively. **(b)** Rotational speed (μm/min) of slow (control stage 4), fast (control stage 7) and not rotating (*fat2* mutant) egg chambers is shown. Red line indicates mean. **(c)** Intracellular MRLC::GFP individual dot-like signals were analyzed at the basal side of follicle cells. Direction of their movement (based on time projected images) was expressed as angle on range of 0°-360°. Example for control stage 7 (fast epithelial rotation) is shown. A = anterior, P = posterior. Scale bars = 5μm. Anterior is on the left. **(d)** Angular distribution of MRLC::GFP movement expressed as frequencies plotted in 20 degree-bin rose diagrams during fast epithelial rotation (stage 6 and stage 8) is shown. Frequencies of MRLC::GFP movement in four 45 degree quadrants are plotted, showing that significant (^***^ = *P*<0.001) majority of MRLC::GFP moves within Up and Down quadrants. Compare with the angular distribution and quadrant frequencies of LifeAct::GFP. Number of analyzed MRLC::GFP and LifeAct::GFP signals are indicated in the lower right with number of analyzed, independent egg chamber (in brackets).

**Extended Data Figure 2.**
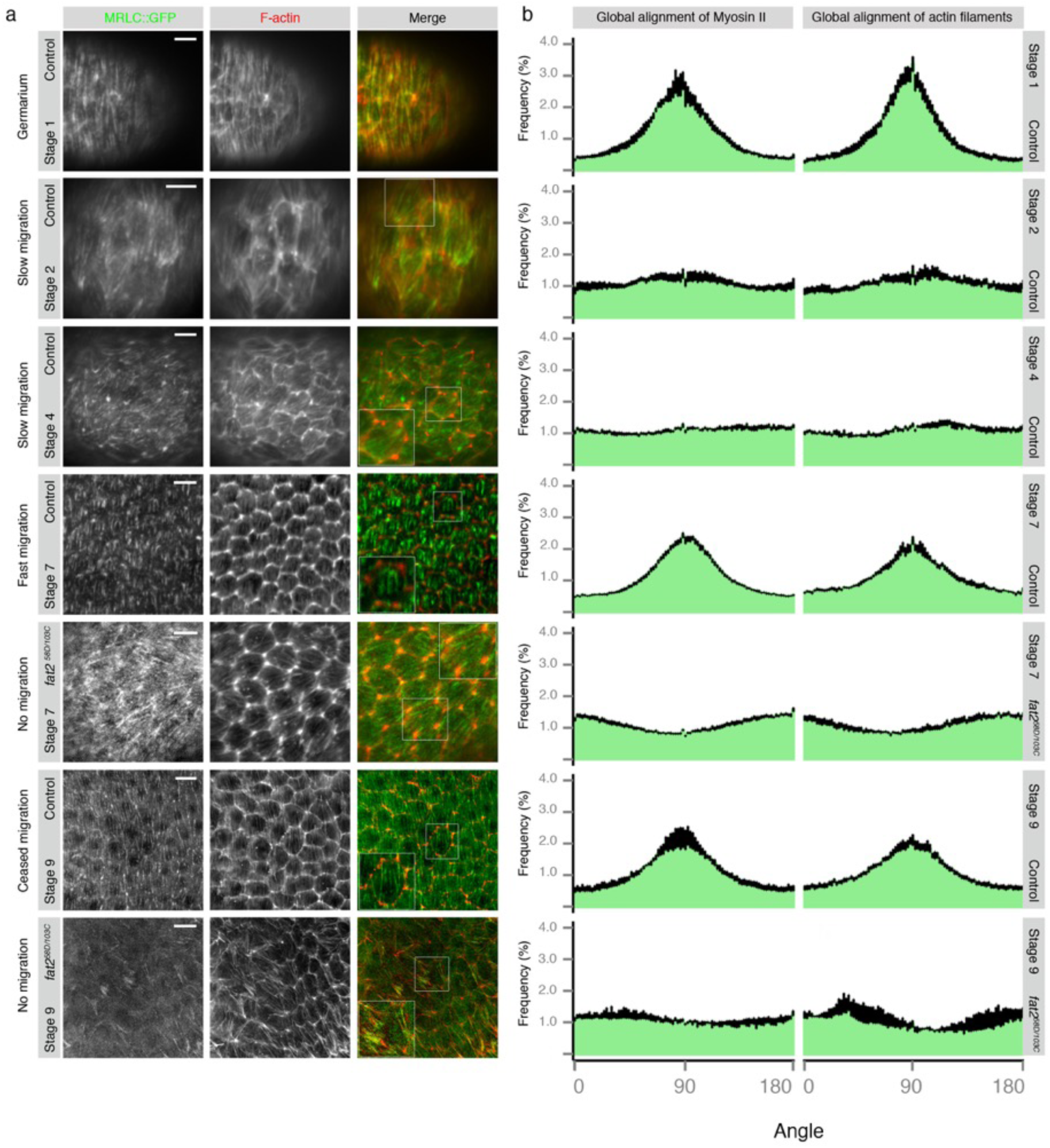
**(a)** Global alignment of MRLC::GFP (green) and actin filaments (red) to the AP axis (0°-180°) at the basal side of the *Drosophila* germarium, showing strong perpendicular alignment to the AP axis, which is temporarily lost when the egg chamber buds from the germarium (stage 2) during early oogenesis (represented by control stage 4) and reaches its proper perpendicular alignment at the time of fast epithelial rotation (represented by stage 7), which is still present at stage 9 when egg chambers cease their epithelial rotation. In *fat2* mutant fixed egg chambers, the MRLC::GFP planar polarized pattern was globally disturbed and reflected direction of actin filaments at stage 7 and stage 9. White boxes show magnification of a representative follicle cell of a particular stage, which display local MRLC::GFP signal localization. Note that MRLC::GFP displays irregular signal distribution in *fat2* mutant egg chambers (stage 9) compared to corresponding controls. Fan-like MRLC::GFP pattern was observed at control stage 4 and *fat2* mutant stage 7. **(b)** Histograms represent frequency distribution of angles of MRLC::GFP and actin filaments measured between 0° and 180°. Anterior (0°) is on the left, posterior (180°) is on the right. Scale bars = 5μm, except of stage 9 where scale bar = 10μm.

**Extended Data Figure 3.**
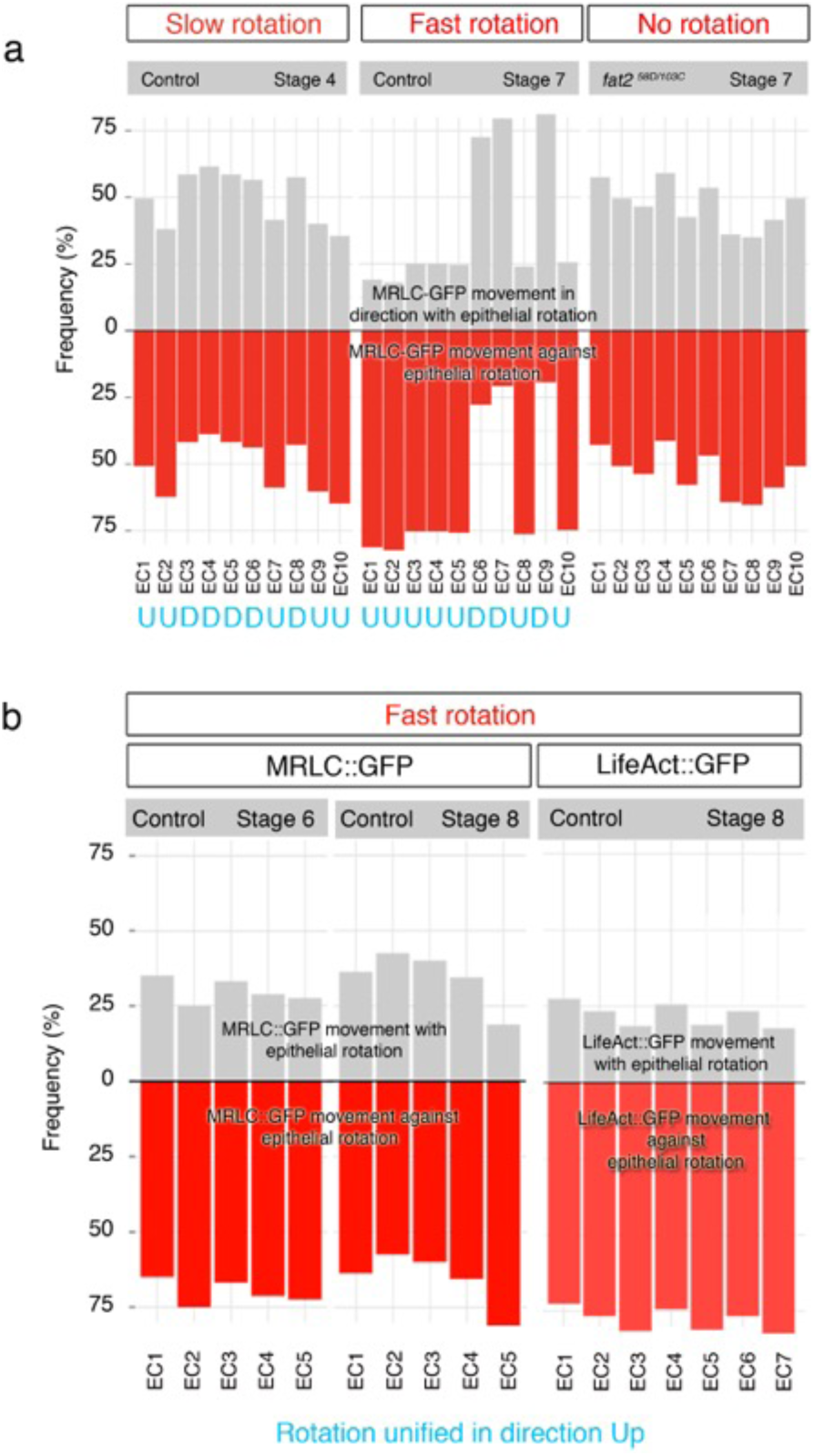
**(a)** Frequencies of MRLC::GFP movement in Up (45°<135°) and Down (225°<315°) quadrants during slow (control stage 4), fast (control stage 7) and no (*fat2* mutant egg chambers) epithelial rotation. Egg chambers were not unified for epithelial rotation. Direction of epithelial rotation is indicated with U (Up) and D (Down) in blue color. Note the MRLC::GFP asymmetry in individual egg chambers that links to direction of epithelial rotation. **(b)** Frequencies of MRLC::GFP movement in Up (45°<135°) and Down (225°<315°) quadrants during fast (control stage 6 and 8) epithelial rotation that was unified to direction Up. Compare with the frequencies of LifeAct::GFP movement during fast epithelial rotation (stage 8). The strong retrograde movement of LifeAct::GFP is not significantly stronger than movement of MRLC::GFP of the same stage (see Extended Data Figure 4).

**Extended Data Figure 4.**
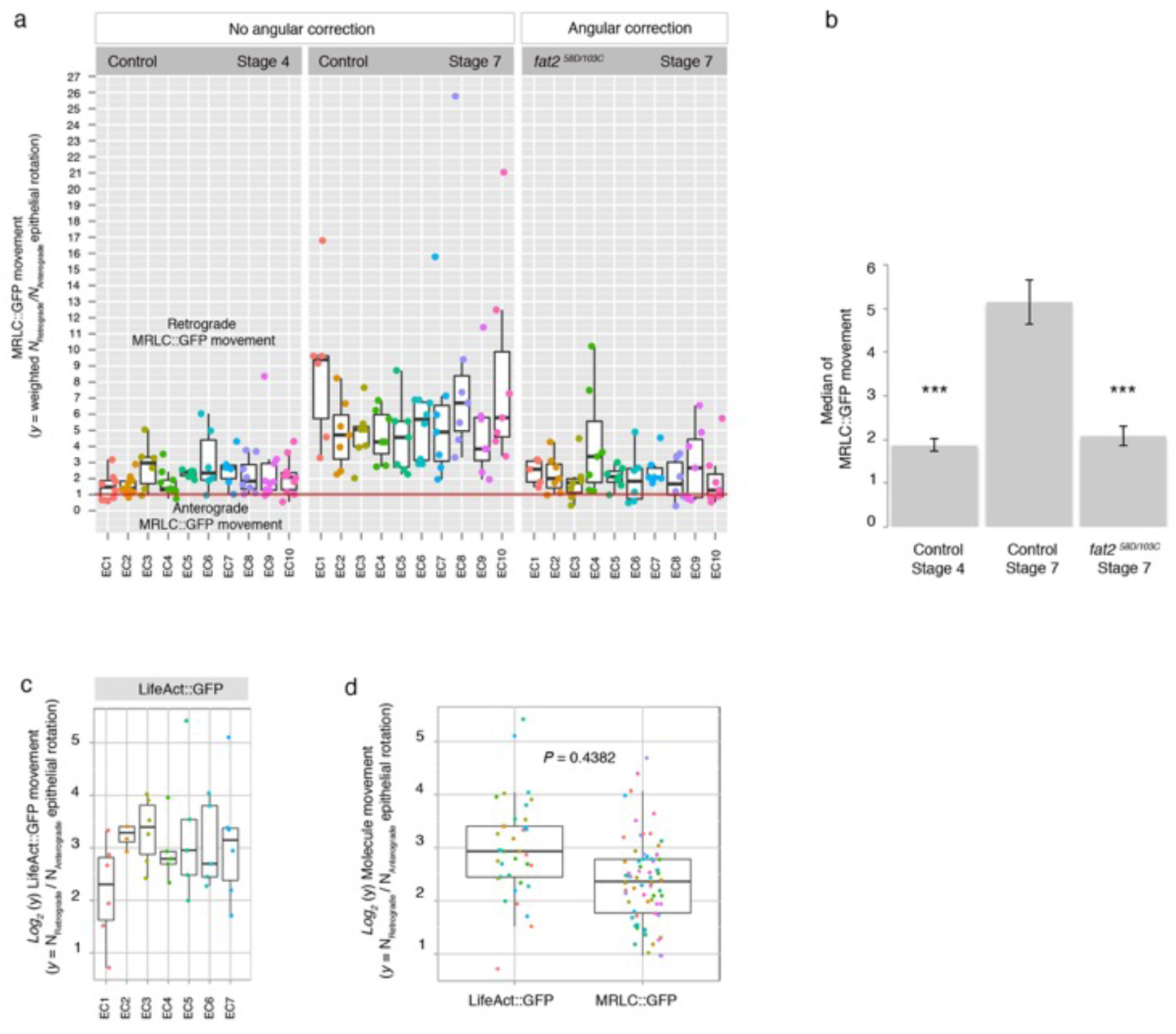
**(a)** Weighted ratios of MRLC::GFP signals moving in the direction Down(225°<315°, retrograde)/Up (45°<135°, anterograde), which are original data corresponding to *log2* scale displayed in Figure 2c. Individual egg chambers (EC) were unified to rotate Up. Symmetry border is indicated with red line. Box plots with medians over all analyzed follicle cells of independent egg chambers with slow (control stage 4), fast (control stage 7) and no (*fat2* mutant of stage 7) epithelial rotation are shown. **(b)** Significantly stronger MRLC::GFP asymmetry [retrograde/anterograde based on **(a)**] expressed as median over all analyzed follicle cells of egg chambers (10 for each type of epithelial rotation) with fast epithelial rotation (control stage 7) compared to slow (control stage 4) and no (*fat2* mutant of stage 7) epithelial rotation. *P*<0.001 (^***^). **(c)** Weighted ratios of LifeAct::GFP signals moving in direction Down(225°<315°, retrograde)/Up (45°<135°, anterograde) for follicle cells of control (stage 8) egg chambers, which were unified to move Up, plotted on *log2* scale show no significant difference, as indicates *P*-value, to *log2* weighted ratios of MRLC::GFP movements **(d)** as shown in Figure 2 and **(a)**.

**Extended Data Figure 5.**
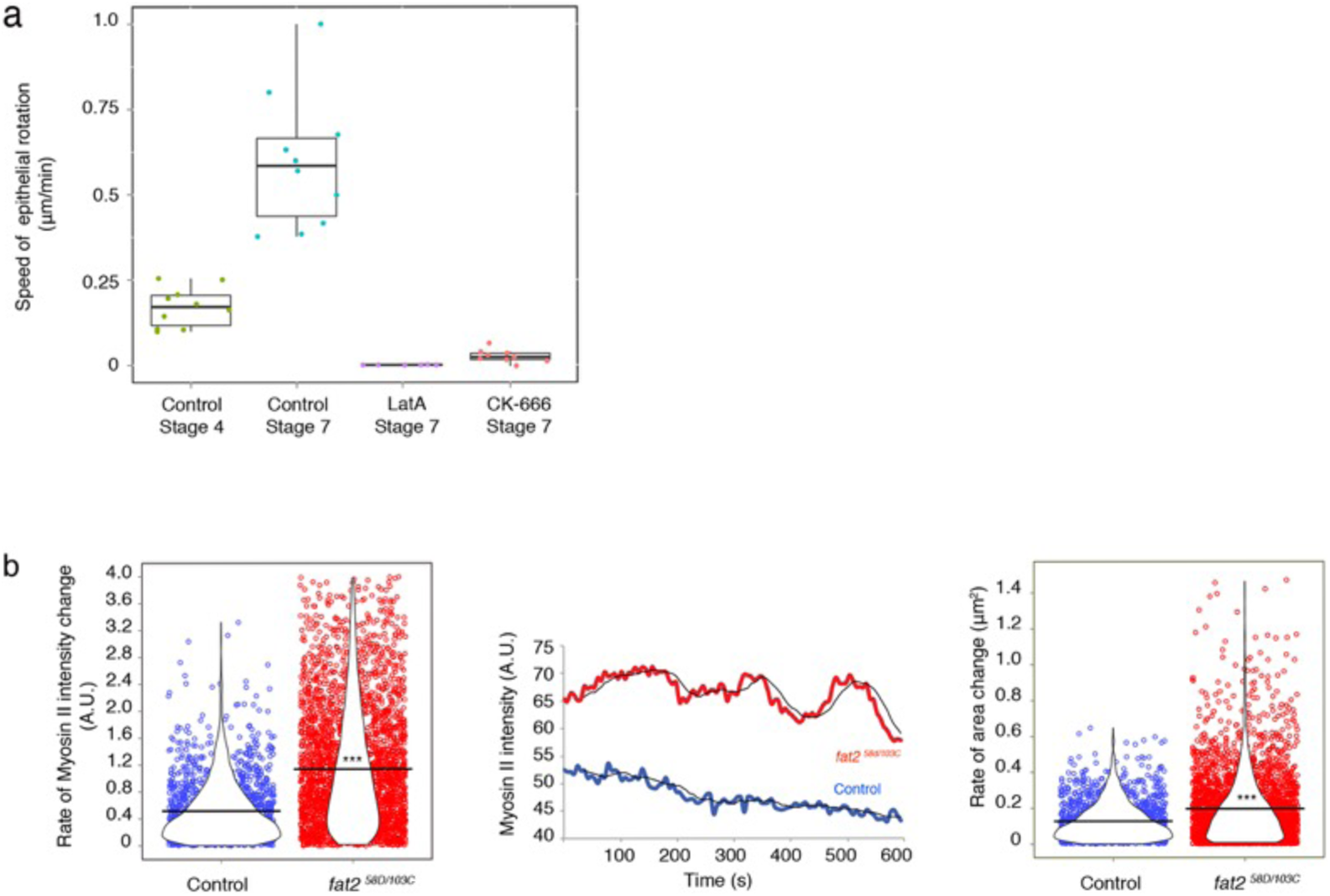
**(a)** Rotational speed of analyzed egg chambers with slow (control stage 4), fast (control stage 7) and no epithelial rotation, which was calculated in egg chambers pharmacologically treated with actin-depleting drug (Latrunculin A) and adhesion-depleting drug (CK-666). **(b)** Rate of Myo-II intensity change (A.U.) and rate of area change μm^2^) when all changes per follicle cells (n = 28, control) over time of 5 independent egg chambers significantly different from *fat2* mutant follicle cells (n = 56) of 7 analyzed *fat2* mutant egg chambers. *P*<0.001 (^***^). Black bars indicate mean. Example comparison of representative follicle cells (control and *fat2* mutant) is shown in original units that measured MRLC::GFP intensity over time (A.U.).

## MOVIES

### Movie 1

Time-lapse movie of MRLC::GFP (green) signals moving at the basal cortex of young slowly rotating egg chamber (control stage 4). Membrane marker stains cell outlines (red). Frame interval = 6s. Scale bar = 5 μm. Anterior is on the left.

### Movie 2

Time-lapse movie of MRLC::GFP (green) signals moving at the basal cortex of mid-oogenesis fast rotating egg chamber (control stage 7). Membrane marker stains cell outlines (red). Note the MRLC::GFP directed movement that is perpendicularly to the AP axis of the egg chamber. Preferred direction against the epithelial rotation was revealed only upon an angular quantification (see Online Methods and Figure 1 and Figure 2). Large MRLC::GFP dots always position towards the lagging side of follicle cells. Frame interval = 6s. Scale bar = 5 μm. Anterior is on the left.

### Movie 3

Time-lapse movie of MRLC::GFP (green) signals moving at the basal cortex of the static *fat2^58D/103C^* mutant mid-oogenesis egg chamber (stage 7). Membrane marker stains cell outlines (red). Note that MRLC::GFP is locally (within a cell) polarized (directed subcellular movement shown in Figure 2) but its global polarity is lost (Figure 1). Large MRLC::GFP dots were also lost. Cell outlines display deformations linked to MRLC::GFP increased intensity (Figure 4). Frame interval = 6s. Scale bar = 5 μm. Anterior is on the left.

### Movie 4

Time-lapse movie of MRLC::GFP (green) signals at the basal cortex of static, actin-depleted (Latrunculin A) mid-oogenesis egg chamber (control stage 7). Membrane marker stains cell outlines (red). Note that MRLC::GFP movement is ceased and MRLC::GFP large dots affected in their position. Frame interval = 6s. Scale bar = 5 μm. Anterior is on the left.

### Movie 5

Time-lapse movie of MRLC::GFP (green) signals at the basal cortex of CK666 (ARrp2/3 inhibitor) treated mid-oogenesis egg chamber of stage 7, which staled the epithelial rotation. Membrane marker stains cell outlines (red). Note that MRLC::GFP signal movement centers as well as MRLC::GFP large dots cross-link each other and position towards the center of follicle cells. Frame interval = 6s. Scale bar = 5 μm. Anterior is on the left.

### Movie 6

Time-lapse movie of *Act5Gal4*>*UAS-LifeAct::GFP* signals at the basal cortex of control stage 7-8. Direction of the epithelial rotation is downwards. Note that the driver used allowed patched expression of *LifeAct::GFP* to identify front (leading edge) actin protrusions of follicle cells. Frame interval = 6s. Scale bar = 5 μm. Anterior is on the left.

## ONLINE METHODS

### Fly stocks and genetics

*Drosophila* MRLC (myosin regulatory light chain of the non-muscle conventional Myosin II), encoded by *spaghetti-squash* (*sqh*), was visualized by MRLC fused with eGFP^27^ under the *sqh* promoter in a null *sqh^AX3^* or *sqh^AX3^/sqh^AX3^;;fat2^58D^/fat2^103C^* mutant background to avoid competition with the endogenous protein. The following stocks: *sqh^AX3^/ sqh^AX3^; sqh-MRLC::GFP/ sqh-MRLC::GFP* (on the II. chromosome) and newly established *sqh^AX3^/sqh^AX3^;sqh-MRLC::GFP/ sqh-MRLC::GFP; fat2^58D^/fat2^103C^,* were used in all figures except of Extended Data Figure 2, where *sqh-MRLC::GFP/ sqh-MRLC::GFP*; *fat2^58D^/fat2^103C^* line was used for *fat2* mutant egg chambers. To visualize actin filaments by time-lapse life imaging, we used *Act5C-Gal4* (BL4414, on II. chromosome)>*UAS-LifeAct::GFP* (on II. chromosome) shown in Movie 6. To label cell membranes we used CellMask^™^ Deep Red (Invitrogen).

### Time lapse imaging

Egg chambers were cultured and life imaging performed as described^9^. An inverted LSM 700 Zeiss confocal microscope was used with 63x/1.45 water immersion lens. Time-lapse movies were taken with an interval of 6s for 300s-600s.

### Fixation and immunohistochemistry

Adult fly ovaries were dissected in 1×PBS and fixed with 4% *p*-formaldehyde for 20minutes. Immunostaining followed standard protocols. We used polyclonal GFP tag antibody conjugated with Alexa Fluor 488 (Molecular Probes) in dilution of 1:100 and rhodamine-phalloidin in dilution of 1:200. Images were acquired on inverted LSM700 Zeiss confocal microscope with 63x/1.45 oil immersion lens.

### Drug treatments

To inhibit polymerization of actin filaments, Latrunculin A (10 μΜ in 1 % DMSO, Enzo Life Sciences) was used for ca. 10mins before direct imaging. To deplete actin protrusions and follicle cell adhesion to the ECM, we used Arp2/3 inhibitor (CK-666, 250 μΜ, Sigma) for ca. 1h as described^19^.

### Image processing, data analysis and statistics

#### Measurement of direction of MRLC::GFP (small dot-like signals) movement

To measure the direction of MRLC::GFP movement in individual follicle cells and globally in the epithelial tissue, we measured angle relatively to the AP axis of egg chambers (Extended Data Figure 1c) with ‘Angle’ tool in Fiji. Before angle measurement, time-lapse movies were corrected for bleaching and cell membranes were registered for their movement to make them static. Angles were then measured on time (60s) projections of MRLC::GFP signals. Altogether, we measured movement of MRLC::GFP relatively to the cell membrane.

To unify *fat2* mutant egg chambers with no epithelial rotation in order to compare with the weak and fast epithelial rotation data, the higher value of MRLC::GFP movement identified for Up (45°<135°) and Down (225°<315°) quadrants was artificially assigned to the Down quadrant to mimic as if all egg chambers rotated upwards, i.e. the preferred MRLC::GFP movement was against (retrograde) epithelial rotation (used in Figure 1c and Figure 2b,c,d).

#### Measurement of Myo-II size and velocity

The size of small and big MRLC::GFP signal was measured as a diameter over 5 and 10 independent egg chambers, respectively. Myo-II velocity was measured on a displacement of Myo-II signals in 30s-60s in original time-lapse movies over 10 independent egg chambers.

#### Measurement of velocity of epithelial rotation

The velocity was defined as an average velocity over 3 independent measurement of cell membrane movement in the most central part of the confocal plane.

#### Angular correction

To define a direction of the MRLC::GFP movement within follicle cells in *fat2* mutant egg chambers, time projected MRLC::GFP pattern served as a definition of the main MRLC::GFP arrays that were reoriented within the existing smallest angle (≤90°) to achieve the perpendicular orientation to the AP axis (used in Figure 2). We hypothesized that Fat2 reorients Myo-II of follicle cells either by spatial regulation of the intracellular actomyosin dynamics or via remodeling of adherens junctions. Thus, in both cases, we concluded that it will be an energy-demanding process and angularly corrected follicle cell perpendicularly to the AP axis of egg chambers (based on the time projected Myo-II pattern) in the smallest possible angle (e.g. as shown in Figure 2a, 45° clockwise and not 135° anti-clockwise).

#### Quantification of global Myo-II and actin filament alignment

To measure global alignment of MRLC::GFP and actin filaments in fixed tissues, we used Fiji software ‘Directionality’ http://imagej.net/Directionality.

#### Measurement of follicle cell shape and elongation direction

Outlines of follicle cells were used to measure Roundness parameter (Shape descriptors>Roundness: 4×[*Area*]/*π*×[*Major axis*]^2^, Uses the heading Round) in Fiji. Angles for definition of direction of elongation of follicle cells (longest axis) were measured with ‘Angle’ tool in Fiji. Angle for definition of the Myo-II direction was defined on time projected MRLC::GFP pattern.

#### Measurement of Myo-II intensity

MRLC::GFP intensity was measured as a mean intensity in the defined cell outline of follicle cells. We developed following pipeline: To measure MRLC::GFP intensity and area of follicle cells, cell segmentation was performed using a custom macro from IRB (Barcelona, Spain) http://adm.irbbarcelona.org/image-j-fiji. Afterwards, cells between subsequent time frames were associated with each other if they overlapped by more than 60 % according to the Jaccard index metric. Determination of mean intensity, cell area and cell roundness was done using an in-house developed plugin utilizing the Analyze Particles Plugin http://imagej.net/ParticleAnalysis in Fiji^41^. Analyses were done on time-lapse movies (6s frame interval), for each frame one plane in z-axis was imaged. Bleach correction was applied in Fiji.

Amplitude of MRLC::GFP intensity change and cell area change was measured in the way that a mean of MRLC::GFP intensity and cell area was plotted over time (360s) of individual cells of a type (control vs. *fat2* mutant), smooth curves were fitted, detrended and a value (based on detrended max.-min. values) calculated for each individual cell of a type and averaged over all cells of the same type.

The time series were smoothened with a Gaussian filter with window of 10 data points (i.e. 1min.) in R Studio http://www.rstudio.com. Rate of MRLC::GFP change was calculated as a first derivative of measured intensity. Rate of basal area reduction was the inverse value of the first derivative of basal area.

Cross-correlation efficiency was calculated with time shift from −6 to +6 min in R Studio. Rate of MRLC::GFP and area was used to calculate the cross-correlation coefficient. The average correlation coefficient was result of averaging smooth correlation data for individual cells.

MRLC::GFP intensity was measured in the most central follicle cells of focal plane of time-lapse movies.

#### Measurement of planar cell chirality (PCC)

One frame of time-lapse movies was selected and the AP axis of egg chambers was turned 90° clockwise or anti-clockwise in order to unify direction of epithelial rotation to the right. Oblique angle (*θ*) was measured for all cell membranes between 0°-90° (right) and −90°-0° (left) range, right and left frequencies were summarized to define to what side (left or right) cell membranes prefer to tilt as described ^6^.

#### Data plotting, statistics and cartoons

Rose diagrams, quadrant plots, histograms, bar and box plots were assembled in R studio http://www.rstudio.com. using various packages ^42 43 44 45^. Error bars represent standard error of the mean (s. e. m.). An unpaired two-sided *t*-test was used. Cartoons were made in Adobe Illustrator CS 5.1.

